# Single-cell characterisation of the hematopoietic bone marrow interactome in health and disease

**DOI:** 10.1101/2022.05.13.491790

**Authors:** Sarah Ennis, Alessandra Conforte, Eimear O’Reilly, Tatiana Cichocka, Sukhraj Pal Dhami, Pamela Nicholson, Philippe Krebs, Pilib Ó Broin, Eva Szegezdi

## Abstract

The bone marrow (BM) is a complex microenvironment and the primary site of hematopoiesis, coordinating the production of billions of blood cells every day. Despite the essential role of the hematopoietic niche in maintaining hemostasis and its relevance to hematopoietic diseases, many aspects of this environment remain poorly characterised due to experimental hurdles. Here we present a high-resolution characterisation of the niche in health and acute myeloid leukemia (AML) by establishing a comprehensive single-cell gene expression database of nearly 340,000 BM constituent cells encompassing all disease stages (healthy BM, AML at diagnosis, remission and relapse). We characterised the cell type composition of the BM and found that the proportions of both myeloid and lymphoid lineage cell types are significantly altered in AML. We also determined broadscale dysregulation of gene expression in almost all BM cell types upon establishment of AML, indicating that the entire niche is disrupted by the disease. Given the importance of interactions between hematopoietic cells and their microenvironment in regulating their function and properties, we determined all possible ligand-receptor interactions between hematopoietic stem and progenitor cells (HSPC) and every other BM constituent cell type. This analysis revealed a remarkable expansion of HSPC interactions in AML involving multiple BM constituent cells that can drive dysregulated HSPC-cell adhesion, immunosuppression and enhanced cytokine signalling. In particular, we found that interactions involving TGFB1 become widespread in AML and present evidence that these interactions can drive AML cell quiescence in vitro, thus highlighting TGFB1 signalling as a potential target for increasing drug sensitivity and preventing relapse. Our results shed light on potential mechanisms of enhanced competitiveness of AML HSPCs and an overall skewed microenvironment that fosters AML growth.

## 2 Introduction

The hematopoietic niche within the human BM consists of a diverse range of hematopoietic and non-hematopoietic (such as endothelial, mesenchymal and osteolineage) cells [1]. The role of the niche is to regulate hemostasis by coordinating the homing, self-renewal, proliferation and differentiation of hematopoietic stem and progenitor cells (HSPCs) and orchestrate the daily production of myeloid, lymphoid and erythroid lineage blood cells. This environment is tightly regulated through reciprocal interactions between HSPCs and the niche [2]. For example, C-X-C motif chemokine ligand 12 (CXCL12)-expressing mesenchymal stromal cells play a crucial role in supporting HSPCs by tethering them in the BM via interaction with CXCR4, thus facilitating HSPC quiescence and cell survival [3, 4]. These regulatory interactions are known to be disrupted in leukemia, where malignant cells alter the niche to support their proliferation and survival. At the same time, these changes make the microenvironment less hospitable to normal HSPCs, leading to their egress and impaired blood cell production [5].

This is also the case in acute myeloid leukemia (AML), an aggressive blood cancer characterised by rapid accumulation of myeloid lineage precursor cells in the BM. Mouse models of AML have shown that the BM-engrafting AML cells gradually erode the stroma leading to an over 90% non-erythroid stromal cell loss in advanced disease [6]. This skewed microenvironment is known to drive drug resistance and consequent relapse [7], but the mechanisms and the responsible cellular interactions are not well understood [8]. Better characterisation of the cell-cell interactions in the BM and how they change in AML may provide opportunities for a “two-pronged” treatment approach by directly targeting the cancer cells with chemotherapy while also disrupting their interactions with the microenvironment so that it becomes inhospitable to malignant cells.

Advancements in technology, such as improved mouse models and the development of single-cell and spatial transcriptomics have increased our understanding of the biology of the normal hematopoietic niche, particularly in mice [9, 10]. However, a large-scale characterisation of the composition and behaviour of the niche in humans and its perturbation in malignancy is lacking. Although single-cell RNA-sequencing (scRNA-seq) has been applied to the study of the BM and has provided valuable insights [9–18], due to economic and experimental hurdles, most of these studies have remained restricted to mouse models or cells from few donors. Also, as these experiments are typically performed on BM aspirates, cells that are more adherent, such as bone marrow stromal cells (BMSC) are under-represented [19]. Another challenge when using scRNA-seq to characterise the leukemic BM is the very high degree of heterogeneity between patients. Because these studies typically only include tens of patients, it is difficult to identify unifying mechanisms of leukemogenesis, allowing only patient-level analysis [13–15].

Here we report a large-scale analysis of the bone marrow cell-cell interactome in health and upon development of AML. By integrating multiple scRNA-seq datasets (Table 1), a large database has been generated (339,381 cells from 63 donors) that enabled a comprehensive analysis of the HSPC-interactome in the hematopoietic niche. Using this dataset we established a baseline reference of healthy cells and then identified changes in cell type composition, gene expression and ligand-receptor interactions of HSPCs that occur during the development of AML. The results highlight the widespread alterations in the BM cellular interactome in AML, especially impacting cell adhesion and cytokine signalling.

**Table 1:** Summary of datasets included in this study.

| Dataset no. | No. of donors |  | No. of samples | No. of cells | Reference |
| --- | --- | --- | --- | --- | --- |
|  | Healthy | AML |  |  |  |
| 1 | 0 | 10 | 28 | 111,347 | Current study |
| 2 | 3 | 0 | 4 | 20,344 | Granja et al., 2019 [11] |
| 3 | 20 | 0 | 25 | 69,264 | Oetjen et al., 2018 [12] |
| 4 | 4 | 5 | 9 | 100,407 | Petti et al., 2019 [13] |
| 5 | 5 | 16 | 41 | 38,019 | van Galen et al., 2019 [14] |
| **Total** | **32** | **31** | **107** | **339,381** |  |

## 3 Results and Discussion

### 3.1 A harmonised single-cell atlas of healthy and AML BM

To generate a single-cell atlas of the BM in health and AML, we performed droplet-based scRNA-seq on BM aspirates from 10 AML patients collected at key stages of disease progression (diagnosis, post-treatment/remission and relapse, clinical information available in Supplementary Table 1) and integrated the data with other, openly available scRNA-seq datasets of healthy and AML BM. We identified 4 suitable datasets [11–14] and processed each of them through a uniform pre-processing pipeline to minimise any study-specific effects (Table 1, and further information on each individual sample is provided in Supplementary Table 2).

To ensure optimal batch correction of the datasets and enable simple reuse and sharing of the atlas, we used scArches (Supplementary Figure 1) [20] to generate a harmonised reference atlas of the BM. The atlas consists of a total of 339,381 cells from 63 donors (Figure 1). Cell types within the dataset were labelled using a combination of predictive and manual annotations. First, cell type labels were predicted for each cell using the dataset published by Granja and colleagues as a reference [11]. We then confirmed the predicted labels and annotated cell types that were not included in the Granja dataset based on expression of well-known marker genes (Figure 1d, 1e). Overall, we identified 19 different cell types including all expected hematopoietic lineage cells, such as hematopoietic stem and progenitor cells, and multiple myeloid, lymphoid and erythroid populations (Figure 1d). Due to the large size of the dataset, we were also able to identify clusters of rare and under-represented cell types such as BM mesenchymal stromal cells and megakaryocytes which could not be identified in any of the individual datasets prior to integration. The integrated dataset can be downloaded from the Zenodo repository accompanying this publication, along with a Google colab notebook demonstrating how to explore and reuse it.

**Figure 1:**
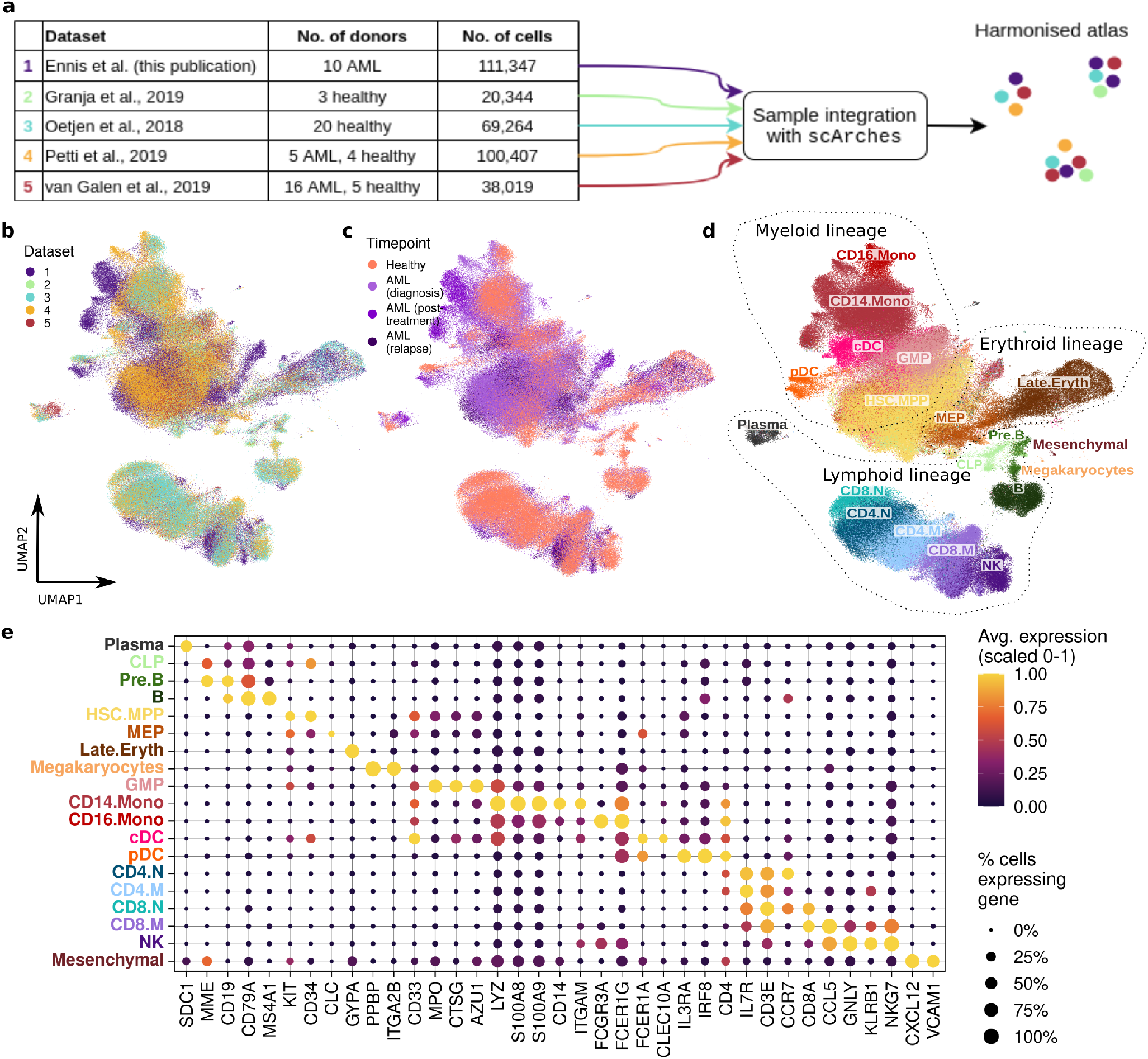
A harmonised single-cell atlas of healthy and AML BM. **a** Schema of dataset generation. **b-d** UMAP was used for dimensionality reduction and visualisation of the dataset. Cells are coloured by dataset (**b**), timepoint (**c**) and cell type (**d**). **e** Dotplot depicting the expression of known marker genes for each identified cell type. The size and colour of each dot represents the percentage of cells expressing the gene and the normalised expression level in each cell type respectively. *HSC*.*MPP = Hematopoietic stem cell/multipotent progenitor, GMP = granulocyte monocyte precursor, MEP = megakaryocyte erythrocyte precursor, CLP = common lymphoid progenitor, cDC = conventional dendritic cell, pDC = plasmacytoid dendritic cell, CD4/8*.*N = CD4/8 naive T, CD4/8*.*M = CD4/8 memory T, NK = natural killer cell*.

### 3.2 Cell type proportions in the BM are significantly altered during the establishment and progression of AML

To understand the HSPC interactome in the BM, first we determined how its cellular composition changes in health and at different stages of AML progression. We used the `propeller` function from the `speckle` R package to test whether there are statistically significant changes in cell type proportions during development and progression of AML [21]. The analysis showed that in healthy BM, over 65% of cells are mature lymphoid lineage cells, around 15% are mature myeloid cells and the rest are progenitor cells, erythroid and other types (Figure 2a). A significant shift in cell type proportions took place at the onset of AML with myeloid progenitor cells making up over 50% of BM cells (Figure 2a). In particular, the most primitive myeloid progenitor population, hematopoietic stem cell/multipotent progenitor cells (HSC.MPP), become significantly enriched at diagnosis. Post-treatment, during remission, these proportions return to levels similar to the healthy condition but at relapse, as expected, enrichment of myeloid progenitors reappears (Figure 2b).

**Figure 2:**
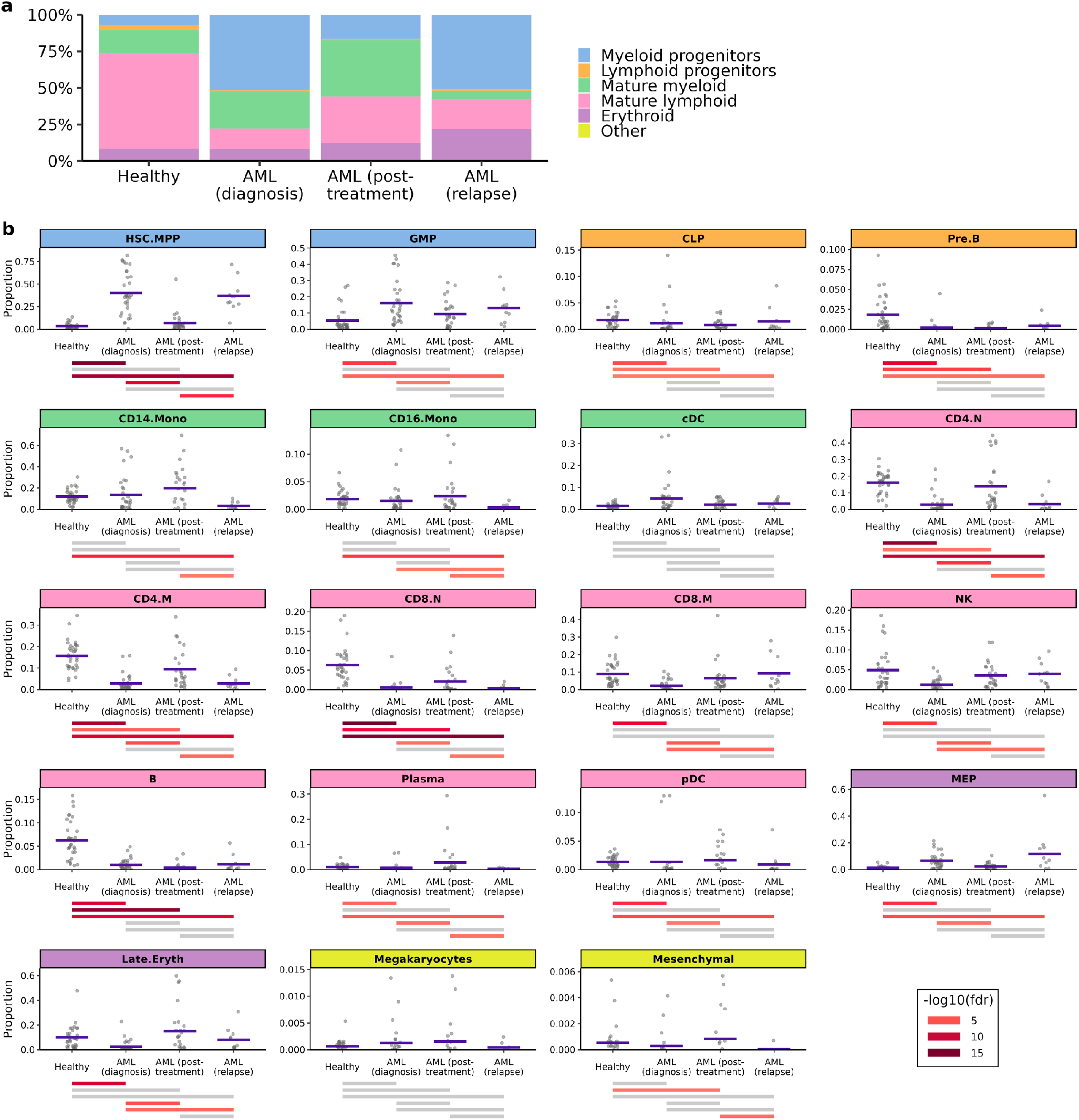
Cell type proportions in the BM are significantly altered during the establishment and progression of AML. **a** Percentage of different cell lineages at each timepoint. **b** The blue bars show the `propeller` estimated proportion of each cell type at different timepoints. The grey dots represent the proportions of each cell type in individual samples. The lines below each plot represent the significance level of the change in proportions between each pair of timepoints. Cell types that showed a significant change in proportions between a given pair of timepoints are represented by coloured lines and grey lines indicate FDR *≥* 0.05. The background colour of the facet labels depicts which lineage (as in **a**) the cell type belongs to. *HSC*.*MPP = Hematopoietic stem cell/multipotent progenitor, GMP = granulocyte monocyte precursor, MEP = megakaryocyte erythrocyte precursor, CLP = common lymphoid progenitor, cDC = conventional dendritic cell, pDC = plasmacytoid dendritic cell, CD4/8*.*N = CD4/8 naive T, CD4/8*.*M = CD4/8 memory T, NK = natural killer cell*.

In addition to these expected changes, profound changes also took place in the lymphoid lineage. The proportion of common lymphoid progenitors (CLP) dropped upon development of AML and failed to return to pre-disease levels in remission. The loss of CLP was associated with a profound drop in its progeny, especially pre-B cells (Pre.B), mature B cells, and to a lesser extent of CD4+ and CD8+ näive T cells (CD4.N, CD8.N) and CD4+ memory T cells (CD4.M). These results reveal that the adaptive immune system suffers long-term damage either due to the disease or to the chemotherapy. Corroborating this notion, B and T cells have been previously reported to display impaired function in AML [22, 23]. This impaired function is likely to be exacerbated by their reduced numbers, which may be important to consider with regard to predicting the efficacy of T- and B-cell-based immunotherapies [24].

A further indication of impaired immune tumour surveillance in AML was that cytotoxic lymphoid cells (CD8+ memory T cells (CD8.M) and natural killer (NK) cells) became enriched at relapse, compared to diagnosis, suggesting that although cytotoxic cell numbers recover during remission, they fail to eliminate the leukemic cells or may in fact play a role in supporting AML cells thus allowing relapse (Figure 2b).

### 3.3 Impact of AML on gene expression in the BM

To uncover functional changes in BM cell types at the onset of AML, we determined changes in gene expression for each cell type upon development of AML. To minimise false discoveries, differentially expressed genes (DEGs) were calculated using a pseudobulk approach, implemented in the Libra R package [25] and the DEGs were then used as input for gene ontology (GO) enrichment analysis. Most cell types showed broad scale changes in gene expression, with a median of 2,368 genes being differentially expressed (|avg. logFC| *≥* 1, adj. p value *≤* 0.05) (Figure 3a). Of all cell types, pre.B, cDC, CD4.M, CD14.Mono and HSC.MPPs had the highest number of DEGs. Notably, pre.B cells showed downregulation of genes mediating B cell maturation (e.g. VPREB1, TCL1A), and upregulated heat shock protein expression (HSPA1A, HSPA1B), indicating cellular stress (Supplementary Figure 2). In HSC.MPPs, genes associated with HSPC self-renewal and differentiation (PBX1 [26, 27]), angiogenesis (VEGFA, NRP1, PDGFA) and myeloid cell differentiation (CSF1R, PTBP3, IL11) were some of the most notable DEGs (Figure 3c). Importantly, the top upregulated gene in HSC.MPP cells, interleukin 1 receptor accessory protein (IL1RAP, avg. logFC = 3.79, adj. p value = 1.94e-33) mediates IL-1*β* signaling, which has been shown to promote the growth and survival of malignant cells in AML and to play a central role in remodelling the BM microenvironment into a niche favoring leukemogenesis over normal hematopoiesis by driving pro-inflammatory signalling [28].

**Figure 3:**
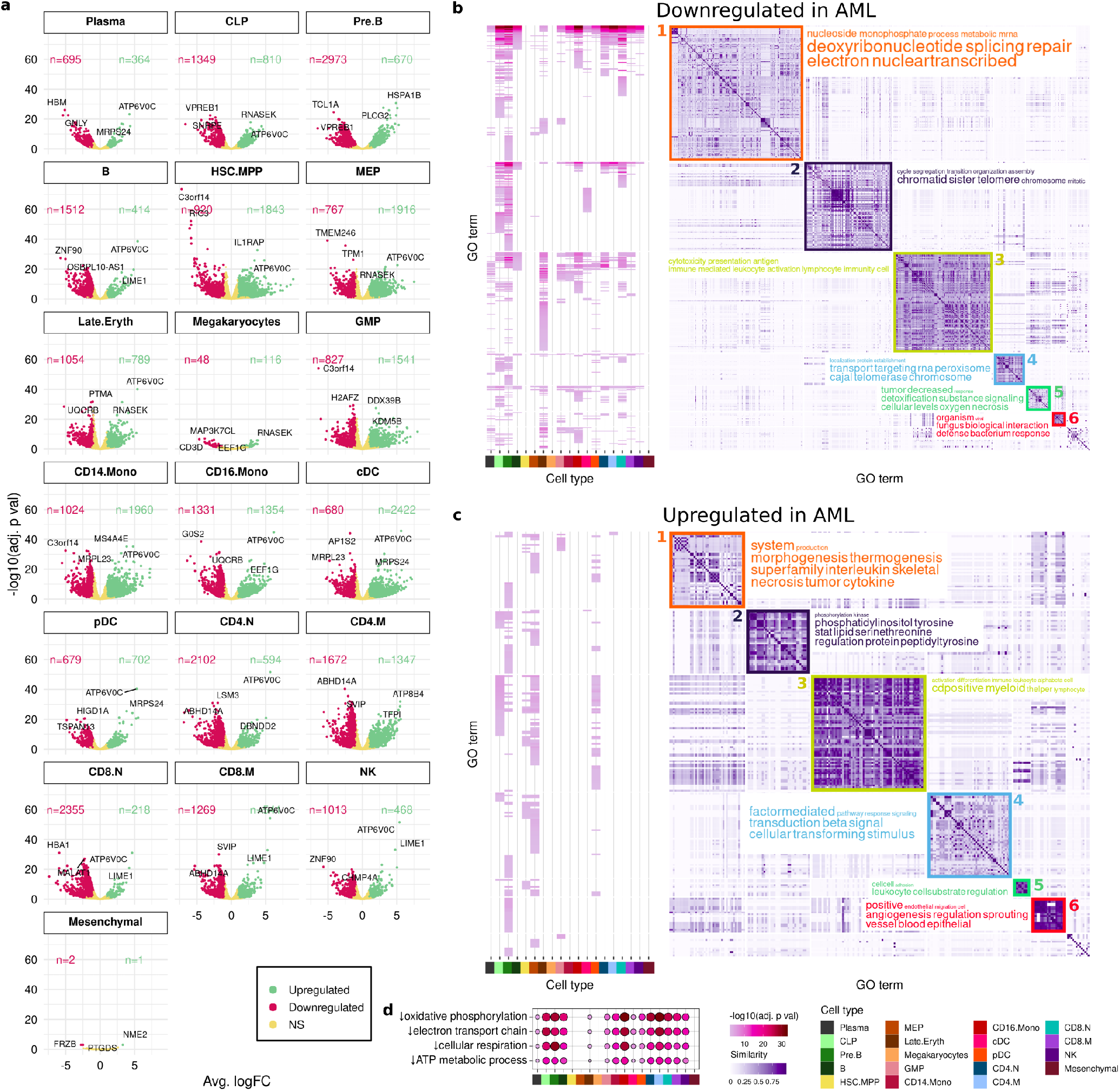
Impact of AML on gene expression in the BM. For each cell type, DEGs were identified by comparing gene expression in AML diagnostic samples to healthy samples. **a** Volcano plots showing number of DEGs for each cell type. **b-c** GO terms enriched among genes significantly down- (**b**) and up-regulated (**c**) in AML. All significantly enriched GO terms are on the vertical axis and the pink heatmap shows the adj. p value for the enrichment of each term in each cell type. The white/purple matrix represents the similarity between GO terms with clusters of similar terms highlighted with coloured boxes. The word clouds beside each cluster show the keywords enriched in that cluster of terms. **d** Enrichment of terms related to metabolism and respiration among genes downregulated in AML. *HSC*.*MPP = Hematopoietic stem cell/multipotent progenitor, GMP = granulocyte monocyte precursor, MEP = megakaryocyte erythrocyte precursor, CLP = common lymphoid progenitor, cDC = conventional dendritic cell, pDC = plasmacytoid dendritic cell, CD4/8*.*N = CD4/8 naive T, CD4/8*.*M = CD4/8 memory T, NK = natural killer cell*.

GO (Biological Process) terms enriched among up- and downregulated genes for each cell type were clustered into groups based on similarity. There were 6 clusters of similar GO terms enriched among downregulated genes and another 6 clusters of GO terms enriched among genes upregulated upon development of AML (Figure 3b 3c). Three clusters of GO terms enriched among downregulated genes affected most BM-resident cell types. These consisted of processes associated with DNA repair, transcription and oxidative metabolism (Cluster 1), lymphocyte functionality (Cluster 3) and oxygen levels (hypoxia), detoxification and necrotic cell death (Cluster 5, Figure 3b). Importantly, the most significantly enriched GO terms among downregulated genes were pathways of mitochondrial respiration (with downregulation of several genes of the mitochondrial electron transport chain: NDUFA4, UQCRB, COX6B1). This was detectable in all BM-constituent cell types, except HSC.MPP, MEP and megakaryocytes, revealing widespread hypoxia and metabolic reprogramming of nearly all non-malignant BM cell types.

Downregulation of genes associated with cell division was also present in multiple cell types, but mostly affected CLP and pre.B cells (Cluster 2). CLP and pre.B cells also displayed impaired mRNA transportation and protein maturation, signs of proteostatic cellular stress (Cluster 4, Figure 3b).

The first group of processes upregulated in AML centred on morphogenesis and cytokine signalling (Cluster 1) predominantly affecting pre.B cells, the malignant population-encompassing HSC.MPP, MEP and GMP populations and pDCs. Of note, there was little overlap in the specific GO terms across these cell types. For example the enriched pathways for MEPs and GMPs differed from each other (e.g. ‘regulation of interleukin-6 production’ enriched among genes upregulated in MEPs but not GMPs).

The second cluster of processes enriched among upregulated genes involved the phosphatidylinositol 3-kinase (PI3K) and signal transducer and activator of transcription (STAT) pathways, especially in CLP and pre.B, where these pathways drive cell proliferation and cell survival before and after immunoglobulin heavy chain and light chain recombination, during which the cells are non-proliferative [29]. Downregulation of mitotic activity (cluster 2 of downregulated pathways) and simultaneous induction of proliferative signalling in these cell cohorts indicates deregulation of B cell receptor assembly (via Ig-recombination) and thus early B cell maturation in AML.

The largest cluster of GO terms enriched among upregulated genes (Cluster 3) was made up of processes related to T cell differentiation, where CLP, Pre.B, HSC.MPP, and pDC cells showed upregulation of genes that drive helper T cell differentiation (IL6R, IL18). This indicates that the leukemic microenvironment may skew lymphoid progenitor maturation towards T cell generation over B cells and may explain the significant B cell depletion observed at all AML timepoints compared to healthy samples (Figure 2b). Other clusters of terms enriched among upregulated genes (Clusters 4 & 5) particularly affected HSC.MPPs and were related to TGF-*β* signalling and cell adhesion pathways.

### 3.4 Interactions between HSPCs and their niche

Cell-cell interactions are central in regulating the hematopoietic BM niche and HSPC maintenance. The role of the niche in controlling HSPC self renewal, expansion and differentiation are well recognised. There is also accumulating evidence that during development of AML, cell-cell interactions change substantially. To gain a comprehensive understanding of HSPC-niche communication pathways and how they change upon development of AML, we predicted ligand-receptor interactions between HSC.MPP and all other BM-resident cell type by looking at the expression of genes that code for ligands and their interacting receptors using the `liana` R package [30]. A combination of interaction prediction methods was run using the Omnipath database of known ligand-receptor interactions [31], which generated a list of high-confidence, well-curated interactions in healthy BM and upon development of AML.

A total of 485 and 905 interactions were identified in healthy and AML diagnostic samples, respectively (Figure 4a). Of the total 1,390 interactions, only 354 were common to both timepoints, demonstrating the extent of the impact that leukemia has on the interactome of hematopoietic cells (Figure 4a). Of the interactions that were common to both timepoints, 34 had a higher interaction score in healthy samples, 43 had a higher score in AML and 277 remained unchanged (Supplementary Table 5, Supplementary Figure 3a). Notably, AML-HSC.MPPs could participate in a large number of interactions which were not present in healthy BM, involving all BM constituent cell types (Figure 4a). This may not be because leukemic HSC.MPPs expressed a higher range of ligands and receptors in comparison to normal HSC.MPPs, instead, it likely reflects the intra-patient heterogeneity of AML where different AML sub-populations co-exist in individual patients and form varied, partially overlapping interactions.

**Figure 4:**
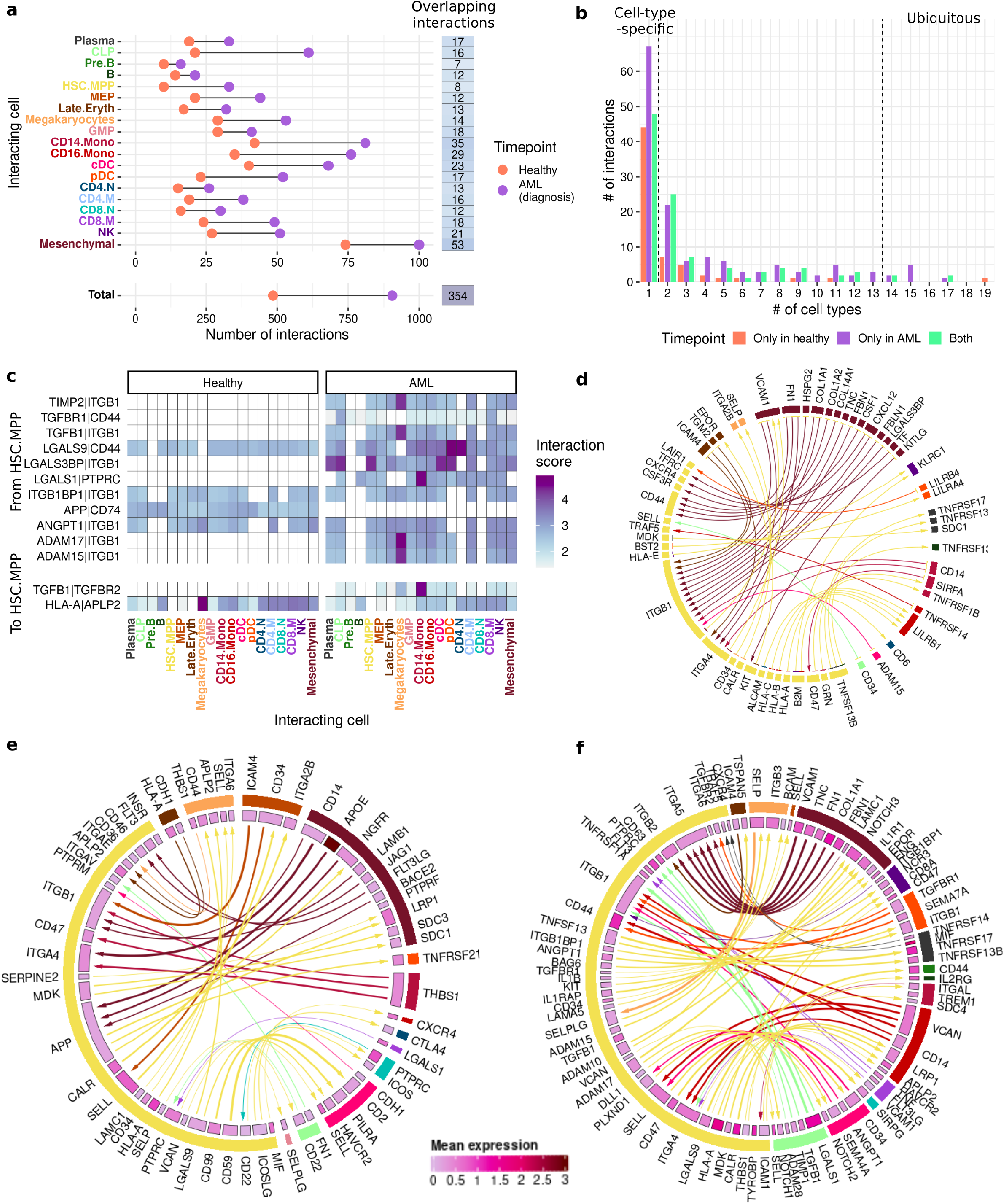
Interactions between HSPCs and their niche. **a** Ligand-receptor interactions identified between HSC.MPP cells and the other cell types in healthy and AML BM. **b** Histogram showing the number of cell types each interaction was identified in. **c** Genes and cell types involved in ubiquitous interactions (*≥*14 cell types). **d-f** Chord diagrams of cell-type-specific interactions identified at both timepoints (**d**), or exclusively in healthy (**e**) or diagnostic (**f**) samples. The colour of the outer sectors represents cell type (colours same as in **a**) and links are coloured by cell type of the sending cell. In **e-f** the inner sectors represent the mean expression of the ligand/receptor gene and the width of links is proportional to the interaction score. *HSC*.*MPP = Hematopoietic stem cell/multipotent progenitor, GMP = granulocyte monocyte precursor, MEP = megakaryocyte erythrocyte precursor, CLP = common lymphoid progenitor, cDC = conventional dendritic cell, pDC = plasmacytoid dendritic cell, CD4/8*.*N = CD4/8 naive T, CD4/8*.*M = CD4/8 memory T, NK = natural killer cell*.

The predicted interactions also varied on a broad scale with regards to how specific they were to given cell types (Figure 4c). We have identified 13 ubiquitous interactions (Figure 4d), where the interacting partner of the HSC.MPP-expressed ligand or receptor was present in at least 14 other cell types (over 75% of BM-constituent cell types). On the other end of the scale, 159 interactions were cell type-specific i.e. the partner ligand or receptor was only present in one cell type (Figure 4e-g). The number of HSC.MPP-interacting cells for the remaining interactions varied, but showed trends where cell types within each hematopoietic cell lineage tended to participate in similar interactions.

The ubiquitous interactions were dominated by cell-adhesion mediators, involving integrin beta 1 (ITGB1), galectins (LGALS1, LGALS9, LGALS3BP) and CD44. Additionally, angiopoietin-1 (ANGPT1) and TGF-*β*1 (TGFB1) signalling was detected (Figure 4d). There was a pronounced shift in these ubiquitous interactions upon development of AML. Firstly, in AML, ITGB1 could form interactions with a broader spectrum of ligands, including the disintegrin and metalloprotease family members, ADAM-15 and -17, as well as the growth factor, angiopoietin-1 (ANGPT1). In addition to enabling adhesion, these interactions could allow HSC.MPPs to receive pro-survival signals from nearly all BM cell types [32]. Another striking change was in the TGFB1-TGFB receptor-2 (TGFBR2) interaction, which became widespread in AML, with CD14.mono cells being a major interacting partner. TGFB1 is a pleiotropic regulator of all stages of hematopoiesis and excessive TGFB1 expression has been linked to proliferation of leukemic cells and failure of normal hematopoiesis in chronic lymphocytic leukemia [33]. In AML, TGFB1 has been linked to leukemia stem cell/leukemia-initiating cell quiescence [4] as well as bone marrow fibrosis [34], however, how TGF-*β*1 signaling changes upon AML and the role of CD14 monocytes in the process has not been understood.

The opposite trend was seen for the interaction between amyloid precursor protein (APP) and the plasma membrane molecular chaperone, CD74, which was widespread in healthy BM, but became almost undetectable in AML. CD74 mediates trafficking of APP into endocytotic vacuoles thus preventing the production of amyloid *β* peptide, the main constituent of senile plaques in Alzheimer’s disease [35]. Loss of the interaction in AML was due to significant downregulation of CD74 expression, which occurred in 13 out of the 19 BM cell types. Loss of CD74 expression may contribute to enhanced fitness of leukemic HSC.MPPs, as CD74 has been reported to regulate HSPC maintenance and its deficiency in mice caused accumulation of HSPCs in the BM due to their increased potential to compete for BM niches [36].

Alternatively, as an MHC-II chaperone, reduced CD74 expression may play an immunosuppressive role, causing reduced MHC-II complex expression, affecting antigen presentation and thus enabling immune escape [37].

With regards to the most active HSC.MPP-interacting cell types, a similar pattern can be seen in healthy and AML BM – bone marrow mesenchymal stromal (BMSC) cells can form the highest number of interactions with HSC.MPPs, followed by CD14+ and CD16+ monocytes (CD14.Mono, CD16.Mono) (Figure 4b). These cell types are known to play important roles in regulating the hematopoietic niche, particularly BMSC, which are essential for HSPC maintenance [2, 38]. The top 50% of interactions between HSC.MPP cells and BMSC and monocytes in healthy BM vs. AML are shown in Supplementary Figure 4.

### 3.5 The hematopoietic stem and progenitor cell and bone marrow stromal cell interactome

Since BMSCs are key regulators of HSPC maintenance, ranging from HSPC quiescence, proliferation, homing and migration to survival, detailed analysis of the BMSC-HSC.MPP interactome was carried out. Both healthy and AML HSC.MPPs formed broad-scale interactions facilitating cell-cell and cell-ECM adhesion with adhesion molecules and ECM components expressed by BMSCs (Figure 5). Interactions between the adhesion receptors, ITGA4, ITGB1 (forming the very late antigen (VLA)-4 complex) and CD44 of HSC.MPPs with collagens, fibronectin, laminins and other BMSC-produced ECM components as well as the cell-cell adhesion ligands VCAM1, VCAN and galectins (LGALS1, LGALS3BP) expressed on the surface of BMSCs were detectable, facilitating HSC.MPP engraftment in the niche. Reciprocally, HSC.MPPs also expressed adhesion ligands, such as galectins (LGALS1, LGALS9), ICAM3 and TIMP1 that could bind to adhesion receptors of BMSCs (CD44, ITGB1, ITGB2, CD63). All these interactions were unchanged, present both in healthy BM and in AML.

**Figure 5:**
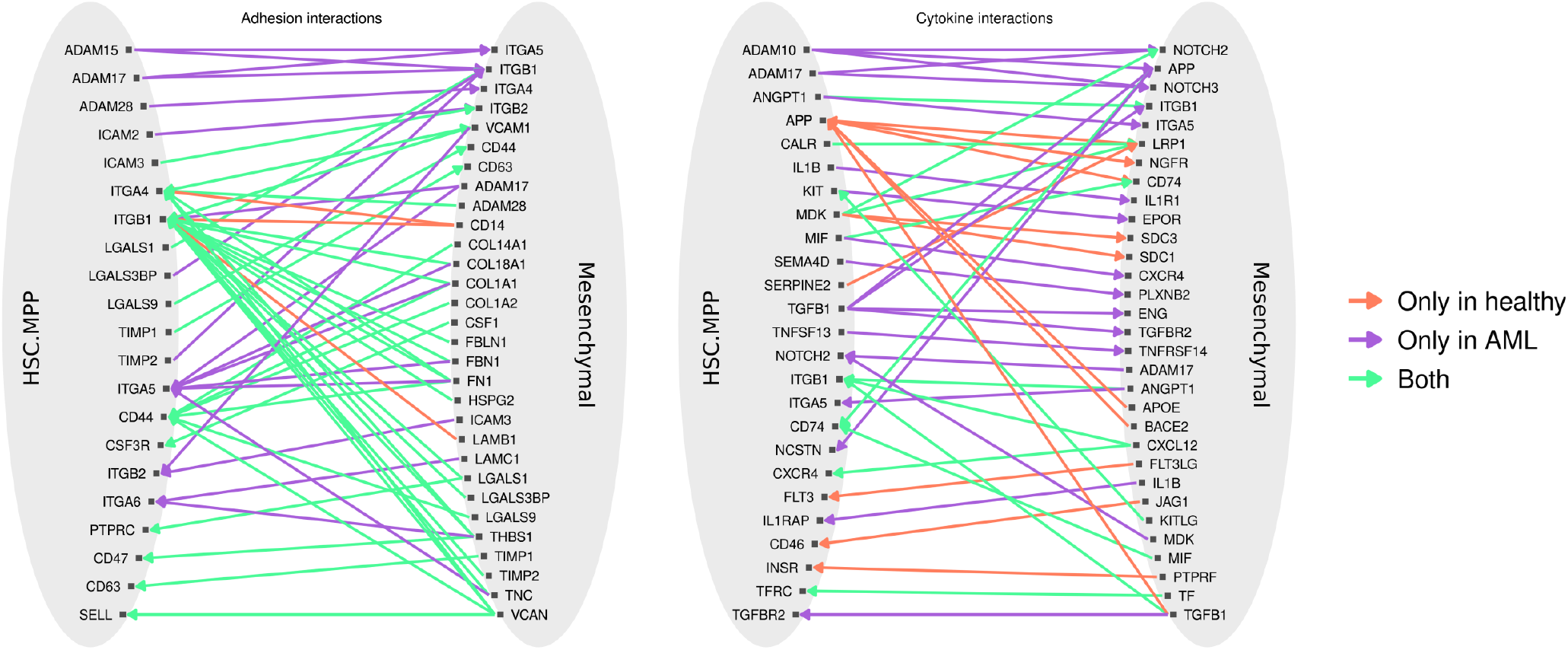
Interactions between HSPCs and BM mesenchymal cells. Interactions were grouped into those involved in cell adhesion and those involved in cytokine signalling. The colour of the arrows represents which timepoint each interaction was identified at. *HSC*.*MPP = Hematopoietic stem cell/multipotent progenitor*

In AML however, a number of additional cell-ECM and cell-cell adhesion interactions became apparent. AML-HSC.MPPs showed increased expression of ITGA5 (avg. logFC = 2.1, adj. p val = 6.42e-14) enabling VLA-5 assembly (ITGA5-ITGB1 complex), induction of ITGA6 (avg. logFC = 1.04, adj. p value = 1.16e-4) as well as of adhesion ligands, most notably members of the ADAM family (ADAM15, - 17). This enabled the formation of further adhesion interactions with ECM proteins (via ITGA5, ITGA6) and BMSC-expressed adhesion proteins (VCAM1 and VCAN via ITGB2), revealing that engraftment of AML-HSC.MPPs in the hematopoietic niche remains essential or it is strongly beneficial to AML cells. Of note, ITGA5 and ITGA6, recognised markers of leukemic stem cells [39], formed interactions with fibronectin (FN) and tenascin C (TNC), an ECM protein important for hematopoiesis and known to co-distribute with FN and collagen III surrounding HSPCs [32, 40].

The expansion of adhesion interactions in AML are likely to increase the competitiveness of AML-HSC.MPPs to engraft in protective niches that provide pro-survival signals and may also contribute to chemoresistance.

In addition to cell-matrix and cell-cell adhesion, BMSCs and HSC.MPPs could engage in a large number of cell-cell and cytokine-cell communication interactions (Figure 5). Many well-established interactions between HSC.MPPs and BMSCs were detected, including CXCL12-CXCR4, KITLG-KIT (Kit ligand and its receptor) and CSF-CSF3R (colony stimulating factor-3/G-CSF and its receptor). In addition, several novel interactions HSC.MPPs formed with BMSCs were also discovered. For example, interactions between MIF (macrophage migration inhibitory factor), a pleiotropic cytokine regulating innate immunity and one of its receptors, CD74; CALR and LRP1 (calreticulin and low density lipoprotein related protein 1), and between MDK (midkine) and two receptors (LRP1 and NOTCH2) were present both in healthy and AML BM. The MIF-CD74 interaction was reciprocal, where both HSC.MPPs and BMSCs expressed both the ligand and the receptor, creating a signaling circuitry driving ERK and PI3K signaling [41].

MDK is a heparin-binding growth factor that can bind to multiple receptors and mediates anti-apoptotic, mitogenic and chemotactic signaling. The interactome analysis found that HSC.MPP-expressed MDK interactions with LRP1 and NOTCH2 were stable, while interactions with syndecan 1 (SDC1) and -3 were lost upon development of AML. On the other hand, a reciprocal interaction became detectable in AML, where AML-BMSC-expressed MDK could bind NOTCH2 on HSC.MPPs, potentiating NOTCH2-signaling in HSC.MPPs. In addition to MDK, AML-BMSCs also induced expression of ADAM17 which can also interact with NOTCH2 facilitating its proteolytic processing and activation. AML-HSC.MPPs thus experience higher NOTCH2-signaling, which has been shown to reduce myeloid differentiation and enhance long- and short-term HSPC repopulation upon stress-hematopoiesis [42]. NOTCH signaling was not only affected in HSC.MPPs, but also in AML BMSCs. ADAM10, 15 and 17 expressed by AML-HSC.MPPs could form interactions with NOTCH2 and NOTCH3 expressed by BMSCs driving NOTCH2/3-activation, which has been shown to block BMSC differentiation and osteogenesis and may enhance BM-mediated drug resistance [43, 44].

In the AML-BM, a number of interactions that were not present in healthy BM also developed. Firstly, IL1B (interleukin-1*β*) produced by HSC.MPPs and BMSCs could activate inflammatory signaling via binding to IL1R1 (IL1 receptor 1) on BMSCs and IL1RAP (IL1 receptor associated protein 1) on HSC.MPPs. Notably, interleukin-1 *β* levels are fine tuned in the BM and chronic IL-1B exposure has been shown to restrict HSPC lineage output and to weaken healthy HSPC self-renewal capacity [45]. IL-1 *β* also promotes AML blast proliferation by induction of growth factors and cytokines like CSF3 [46].

Finally, an interaction between the semaphorin family member, SEMA4D expressed by HSC.MPPs and its receptor, PLXNB2 on BMSCs became apparent in AML. SEMA4D is known to inhibit osteoblast differentiation in multiple myeloma patients, where bone resorption is abundant. This process may be aggravated by TGFB1 signaling, which also became enhanced in AML, with both HSC.MPPS and BMSCs expressed TGFB1 and could activate TGFBR2 on the other cell type. While the effects of TGFB1 on BMSCs are multifold and not fully understood, TGFB1 may reduce their osteoblastic differentiation and facilitate the development of a myelodysplastic syndrome (MDS)/AML-like BMSC phenotype [47].

### 3.6 Altered cell adhesion and cytokine signalling in AML

To determine how the identified interactions regulate cell-cell communication in the BM and thus control the functioning of the hematopoietic niche, ligand and receptor genes were linked with KEGG pathway annotations and filtered to only include interactions where both the ligand and the receptor had the same annotation (Figure 6a). The identified pathways centred around four processes: 1. cell adhesion and migration (including *Cell adhesion molecules CAMS, ECM receptor interaction, Focal adhesion, Adherens junction* and *Leukocyte transendothelial migration*), 2. hematopoiesis and leukocyte maturation (including *Hematopoietic cell lineage, NOTCH signaling pathway* and *Antigen processing and presentation*), 3. cytokine and kinase signalling (*Cytokine cytokine receptor interaction, TGF beta signaling pathway, JAK-STAT signaling pathway, MAPK signaling pathway, Insulin signaling pathway* and *Adipocytokine signaling pathway*) and finally 4. cancer-related pathways (including *Pathways in cancer, Chronic myeloid leukemia, Colorectal cancer* and *Pancreatic cancer*) (Figure 6a).

**Figure 6:**
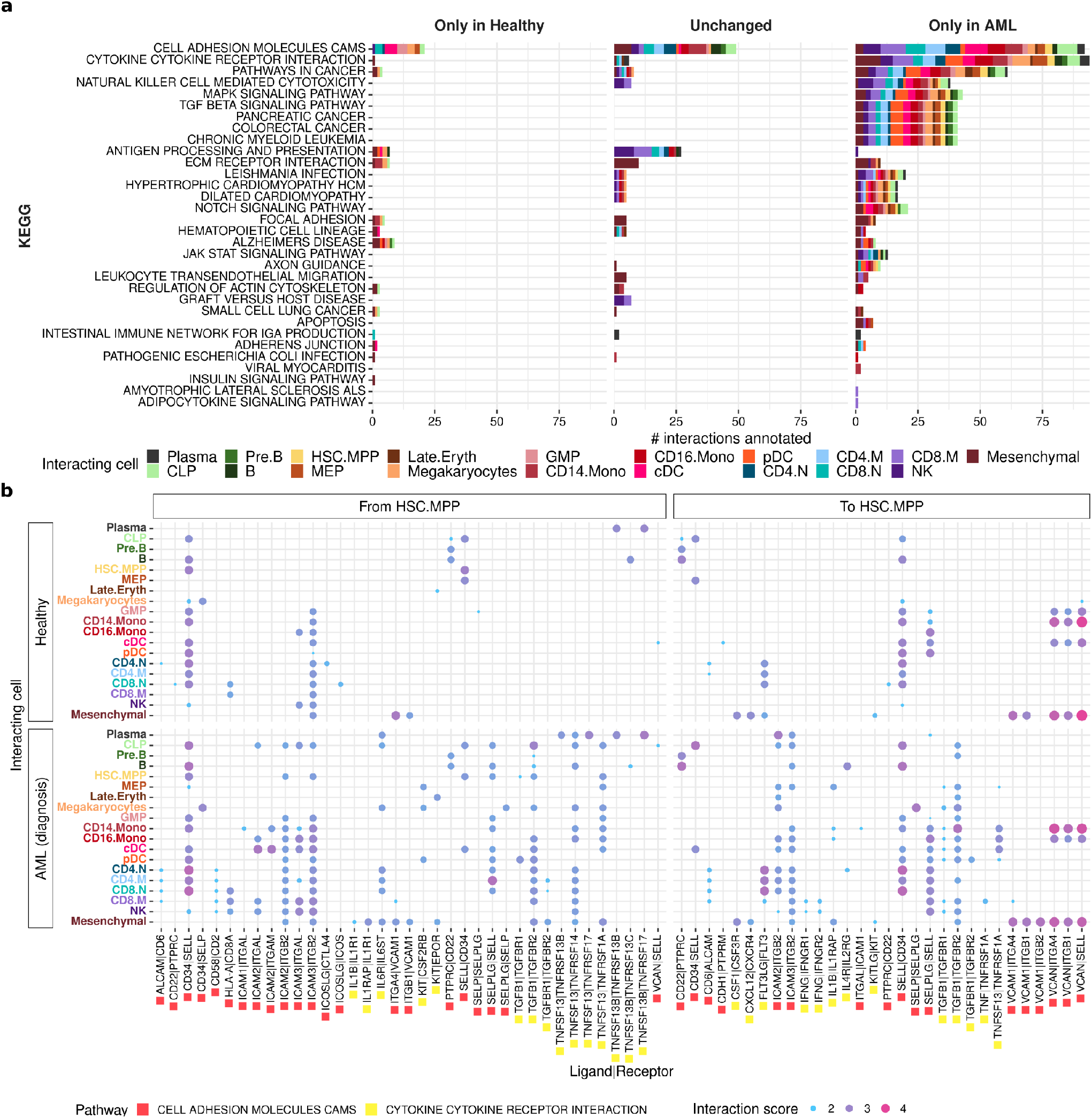
Altered cell adhesion and cytokine signalling in AML. **a** Functional annotation of interactions. Interactions were annotated with KEGG pathway annotations and filtered to interactions where the ligand and receptor gene both had the same annotation. **b** Cell types and signalling molecules involved in interactions annotated with *“CELL ADHESION MOLECULES CAMS”* or *“CYTOKINE CYTOKINE RECEPTOR INTERACTION”*. The size and colour of each dot represents the interaction score which is a measure of the confidence with which that interaction was identified at a given timepoint. *HSC*.*MPP = Hematopoietic stem cell/multipotent progenitor, GMP = granulocyte monocyte precursor, MEP = megakaryocyte erythrocyte precursor, CLP = common lymphoid progenitor, cDC = conventional dendritic cell, pDC = plasmacytoid dendritic cell, CD4/8*.*N = CD4/8 naive T, CD4/8*.*M = CD4/8 memory T, NK = natural killer cell*.

The pathways that the identified ligand-receptor interactions could mediate were restricted and specific to a given cell type/cell lineage in healthy BM (detected only in healthy BM or did not change upon development of AML). Upon development of AML, only a small number of these interactions were lost, instead an expansion both in the number of possible interactions and participating cell types were detected. Additionally, via newly formed interactions, pathways which were not detected in healthy BM were activated, predominantly regulating cancer-related pathways, cell adhesion and cytokine signaling (Figure 6a).

To gain insight into the impact of these interaction on HSC.MPP function and the BM microenvironment in AML, the participating ligand-receptor pairs were analysed for the cell type origin of the signal, the target of the signal if it originated from HSC.MPPs (Figure 6b) and how the interaction changed between healthy and AML BM (Figure 6b).

Adhesion of AML cells to the BM niche can confer protection from apoptosis and enhance chemoresistance and targeting cell adhesion molecules to mobilise malignant cells from the BM has been shown to improve drug sensitivity in AML (e.g. CXCR4 inhibitors) [48]. Corroborating this notion, a broad spectrum of interactions mediating cell adhesion were detected, involving nearly all cell adhesion molecule (CAM) families, namely the immunoglobulin superfamily of CAMs (with members ICAMs, VCAM, ALCAM), integrins (ITGAL (forming LFA-1 when binding to ITGB2), ITGAM (forming Mac-1 with ITGB2) and ITGB2), C-type lectin-like domain proteins (SELL, SELP) and proteoglycans (SELPLG, VCAN) (Figure 6b).

Importantly, cell adhesion interactions between selectin genes (SELL, SELP) and CD34 and the proteoglycan selectin counter-receptor, SELPLG (selectin P ligand/PSGL1) were either upregulated or identified exclusively in AML samples. SELE is typically expressed by endothelial cells and is known to interact with SELPLG expressed on the surface of leukocytes, including AML blasts. Inhibiting SELE and thus preventing AML cell-endothelium/BM interactions (GMI-1271, uproleselan) has been shown to improve AML patient response to therapy [49, 50]. Interestingly, the interactome analysis uncovered another selectin interaction not reported in AML before; between SELPLG (expressed by HSC.MPP cells) and SELP (expressed by megakaryocytes, Figure 6b). The therapeutic potential of SELL and SELP in AML is only partially understood but is supported by recent studies in multiple myeloma that showed that platelets can “cloak” malignant cells via SELP to protect them from NK cell-mediated cytotoxicity [51], and high soluble SELL expression has been found to be associated with AML relapse [52].

HSC.MPPs also participated in numerous interactions involving the integrin family, especially the leukocyte integrins, ITGAL, ITGAM and ITGB2. HSC.MPPs expressed both the integrin ligands, ICAM1-3 and ITGB2, enabling reciprocal interactions between HSC.MPPs and multiple BM-constituent cells. Notably, these interactions substantially expanded in AML. For example, while no interactions were detected for HSPC-expressed ICAM2 in healthy BM, ICAM2-expressing HSC.MPPs were detected in AML and could form cell-cell interactions with 13 out of the 19 BM cell types expressing ITGB2. Interactions via HSC.MPP-expressed ITGB2, ICAM2 and ICAM3 also became widespread in AML, involving 15 cell types (Figure 6b). This expansion of integrin signalling is likely to play a role in promoting survival of the malignant cells and although targeting integrins is challenging, these results warrant the study of new, subtype-specific integrin inhibitors for the treatment of AML [53, 54].

With regards to cytokine signalling, a number of novel as well as previously described interactions were identified. One of the most significant of these was IL1B signalling, which has been shown to promote the growth and survival of malignant cells in AML [28]. Ligands and receptors involved in IL1B signalling (IL1B, IL1R1, IL1RAP) were broadly expressed in HSC.MPP cells, monocytes (CD14.Mono, CD16.Mono) and BMSCs (Figure 6b), corroborating the prevailing inflammatory signalling in AML-BM.

The most outstanding novel cytokine pair interactions highlighted by the analysis was TNFSF13 (A Proliferation-Inducing Ligand, APRIL) and TNFSF13B (B cell activating factor of the TNF-family, BAFF), two key regulators of B cell maturation and survival [55]. TNFSF13B/BAFF is produced by myeloid cells, and exerts its effect by binding to three receptors, TNFRSF13B (Transmembrane Activator And CAML Interactor, TACI), TNFRSF13C (BAFF-R) and TNFRSF17 (B Cell Maturation Antigen, BCMA) [55]. Interaction between TNFSF13B and its three receptors were detectable between HSC.MPPs and plasma cells or B cells in healthy BM and were retained in AML. On the other hand, our analysis identified significant upregulation of APRIL in AML in 13 BM cell types, particularly myeloid lineage cells. APRIL, similar to BAFF/TNFSF13B can also bind to TNFRSF13B and TNFRSF17, while TNFRSF13C is a BAFF-specific receptor. Interaction between TNFSF13/APRIL and its two receptors became apparent in AML-BM corroborating reports that APRIL expression is enhanced in AML cells where it may support leukemic cell proliferation [56]. Underlining the importance of APRIL in malignant transformation, high APRIL expression is also associated with chemoresistance [57] and immunosuppression via facilitating the formation of regulatory B and T cells [58].

The analysis also identified that TGFB1 interactions between HSC.MPP and multiple different cell types was present in AML, while TGFB1 interactions were restricted in healthy BM. As mentioned above, widespread TGFB signaling correlated with upregulation of the two TGF-*β* receptors, TGF*β*-RI (TGFBR1, avg. logFC = 1.27, adj, p value = 1.54e-7) and TGF*β*-RII (TGFBR2, avg. logFC = 1.15, adj. p value = 9.75e-9) - in AML-HSC.MPPs. TGFB signalling is an important regulator of hematopoiesis and has pleiotropic effects depending on the context [59]. In healthy HSPCs, binding of TGFB1 to TGFBR1 and TGFBR2 represses cell proliferation via induction of the cyclin-dependent kinase inhibitor CDKN1C [60, 61]. On the other hand, the response of AML HSPCs to TGFB1 is less well understood [62]. Our data suggest that TGFB signalling is particularly active in AML and the potential of TGFB signalling as a therapeutic target warrants further research.

### 3.7 TGFB1 interactions drive stem cell quiescence in AML

Given the increased prevalence of interactions involving TGFB1 in AML and the upregulation of TGFBR2 in HSC.MPP cells, we analysed the downstream consequences of these interactions.

In the canonical TGF*β* signalling pathway (Figure 7a), TGFB1 binds TGFBR2 leading to the recruitment of TGFBR1, activation of the Smad transcription factors, SMAD2,SMAD3 and transcription of target genes, including cyclin dependent kinase inhibitors thus inhibiting cell cycling. In healthy HSPCs as well as leukemia stem cells (LSC)/leukemia-initiating cells, binding of TGFBR2 activation has been shown to drive quiescence [4, 60, 61], which is linked to drug resistance and new results indicate that inhibition of TGFB1 signalling may enhance the efficacy of chemotherapy [63]. Given these results, we decided to look for evidence of TGFB1-induced quiescence in AML-HSC.MPPs.

**Figure 7:**
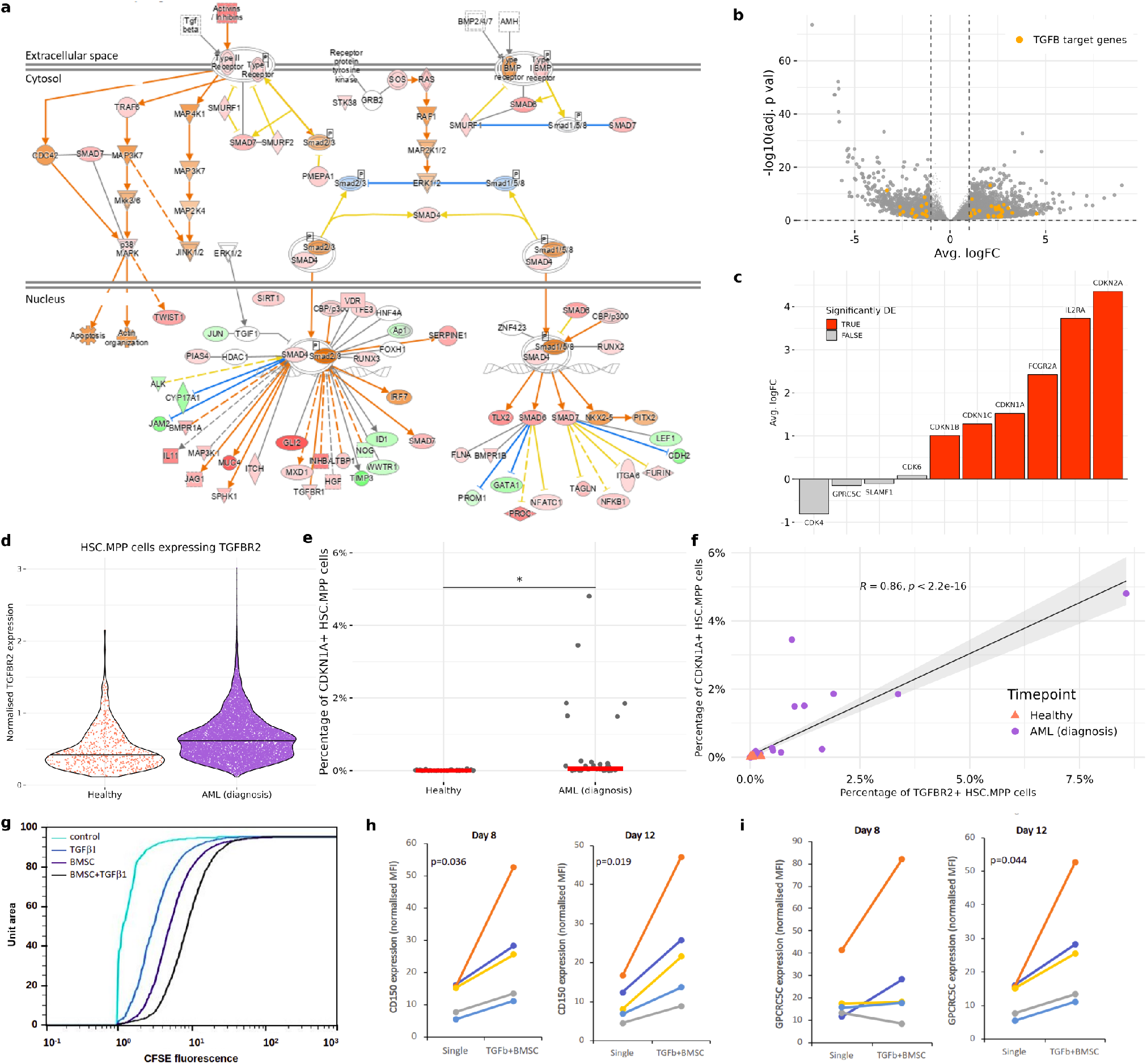
Interactions involving TGFB1 drive stem cell quiescence in AML. **a** The canonical TGFB signalling pathway. Red shaded nodes represent genes upregulated in AML. Orange edges represent predicted upregulation/activation based on canonical pathway. Green nodes represent genes downregulated in AML. Blue edges represent predicted downregulation/inhibition based on canonical pathway. Yellow edges represent interactions opposite to prediction. **b** Volcano plot showing differential expression of TGFB1 target genes in HSC.MPP cells in AML. **c** Differential expression of genes related to HSC/LSC quiescence in HSC.MPP cells in AML. **d** Normalised TGFBR2 expression in HSC.MPP cells (only non-zero values shown). **e** Percentage of CDKN1A+ HSC.MPP cells in each sample (CDKN1A+ defined as normalised CDKN1A *≥* 0.2). **f** Correlation between the percentage of CDKN1A+ and TGFBR2+ HSC.MPP cells in each sample (TGFBR2+ defined as normalised TGFBR2 *≥* 0.175). **g** Proliferation (measured by CFSE fluorescence) of KG1a AML cells cultured alone (control), in the presence of 5 ng/ml transforming growth factor beta 1 (TGFb1), bone marrow stromal cell feeder layer (BMSC) or both (BMSC+TGFb1) for 8 days. Surface expression of CD150 protein **h** and GPRC5C **i** determined with immunofluorescence and flow cytometry in primary AML blasts from 5 different donors cultured either alone or in the presence of TGFb1 on BMSC feeder layer. *HSC*.*MPP = Hematopoietic stem cell/multipotent progenitor, BMSC = Bone marrow stromal cell*

First, activation of TGFB1 signaling in AML-HSC.MPPs was confirmed by determining the overlap of genes differentially expressed in AML-HSC.MPPs and genes in the canonical TGFB1 signalling pathway using Ingenuity Pathway Analysis (IPA, Qiagen). TGFB1 was the most significant activated upstream regulator in HSC.MPPs (Figure 7a,b). The differentially expressed genes also showed presence of a gene signature indicating quiescence in the AML-HSC.MPP population (Figure 7c, gene markers of HSC/LSC quiesence were identified from literature). Out of a total of 10 genes, 6 were significantly upregulated, indicating enhanced quiescence in the AML-HSC.MPP population in comparison to healthy HSC.MPPs (Figure 7c). Furthermore, we found increased expression of TGFBR2 and the LSC quiescence marker, CDKN1A in AML-HSC.MPPs (Figure 7d,e) and a significant positive correlation (*R* = 0.86, p *<* 2.2e-16) between the percentage of CDKN1A+ and TGFBR2+ HSC.MPP cells in each sample. This indicates that the increased prevalence of quiescent cells is linked to TGFB1 signalling (Figure 7f).

The induction of stem cell quiescence by TGFB1 in AML cells was also assessed in the low differentiation-status (LSC-like) KG1a AML cell line and in primary patient samples. Exposure of KG1a cells to TGFb1 either as a single agent, or in the presence of bone marrow stromal cell co-culture, slowed their proliferation (measured with CFSE dye retention assay). Similarly, TGFb1 induced the expression of the HSC quiescence markers, CD150 and GPRC5C in primary, patient-derived AML blasts (Figure 7g,h,i). Taken together, these results emphasise the role of TGFB1 in driving AML cell quiescence.

## 4 Conclusions

It is well-established that reciprocal interactions between hematopoietic cells and their niche are essential in regulating their properties and functions, and that these interactions are skewed in AML generating a microenvironment that fosters leukemogenesis. In this study, we provide several advances by determining the interactome of hematopoietic cells in health and AML. By compiling the largest collection of single-cell gene expression data of the human BM to date, we were able to robustly characterise this complex environment. Our study leverages several previously generated datasets [11–14], along with a novel dataset to perform a comprehensive analysis of the hematopoietic niche.

Using this dataset, we determined changes in cell type proportions, cell-type-specific gene expression and putative cell-cell interactions HSC.MPP form with the other BM constituent cell types. The results highlight several recurring themes that provide insights into how the microenvironment is corrupted by AML. Firstly, B cell maturation is permanently damaged in AML, shown by the clear decline in the number of CLP and their progeny, including pre.B and B cells, at all AML timepoints. Unlike T cells, whose numbers partially recover during remission, the significant depletion of B cells persists post-treatment, which likely has implications for the functioning of the adaptive immune system in AML patients even during disease remission. Gene expression changes indicated skewed lymphoid lineage maturation towards T cells rather than B cells and deregulated early B cell maturation, shedding light on the potential reasons for B cell loss in AML.

The predicted HSC.MPP interactions revealed key cell-cell communication pathways that can provide a competitive advantage to AML cells over normal hematopoietic cells including both well-established mechanisms (e.g. CXCL12-CXCR4, FLT3LG-FLT, KITLG-KIT) and several novel ones (SELPLG-SELP, SEMA4D-PLXNB, MIF-CD74, MDK and its receptors, etc). These interactions can drive:

1. Enhanced engraftment of AML cells in protective niches, via enhanced cell-adhesion mediated via integrins (VLA-5, LFA-1, Mac-1), CD44, selectins as well as ADAMs.
2. Immunosuppression, for example by reduced antigen presentation, and the cytokine, APRIL.
3. Impaired BMSC functions/bone resorption via enhanced Notch, TGFB1 and SEMA4D signalling.
4. Establishment of an inflammatory environment (IL1B and IL6).
5. Quiescence of AML cells, mediated by TGF*β* signalling.

While inferring ligand-receptor interactions between cells from gene expression data comes with certain limitations as surface protein expression is not always reflected at the level of mRNA and it lacks the spatial dimension (adjacency of potentially interacting cells), many of the identified interactions are novel and represent potential therapeutic targets to disrupt AML-niche interactions, and warrant further investigation and validation.

In conclusion, this study provides the first single cell-level characterisation of the interactome of HSPCs in the hematopoietic BM niche in health and AML, discovering novel, AML-specific HSPC interactions, and the BM constituent cell source of these signals at a high granularity. The results reveal the scale of niche alterations that take place in AML and identify promising therapeutic targets to overcome niche-mediated drug resistance.

## 5 Acknowledgements

This research was funded by Science Foundation Ireland through the SFI Centre for Research Training in Genomics Data Science (18/CRT/6214), the Horizon2020 INTEGRATE MSCA-COFUND programme (H2020-MSCA-COFUND-945385), Blood Cancer Network Ireland (14-ICS-B3042), DISCOVER-RISE Programme (H2020-RISE-777995) and the Finnish Biobank. We would also like to thank Carsten Riether for his scientific feedback.

## 6 Methods

### 6.1 Datasets

#### 6.1.1 Generation of scRNA-seq dataset

Longitudinally-collected BM aspirates were obtained for 10 AML patients from the time of diagnosis, post-treatment/remission and at relapse from The Finnish Haematology Registry and Clinical Biobank. The mononuclear cell fraction was isolated using Ficoll density gradient separation. Red cell lysis and dead cell removal (Miltenyi Biotec) was carried out prior to generation of single cell droplets with 10X Genomics Chromium Controller. Cell counting and viability assessments were conducted using To-Pro-3 viability dye. Thereafter, GEM generation & barcoding, reverse transcription, cDNA amplification and 3’ gene expression library generation steps were all performed according to the Chromium Next GEM Single Cell 3’ Reagent Kits v3.1 (Single Index) User Guide (10x Genomics CG000204 Rev D) with all stipulated 10x Genomics reagents. Generally, 16.5 *µ*L of each cell suspension (1,000 cells/*µ*L) and 26.5 *µ*L of nuclease-free water were used for a targeted cell recovery of 10,000 cells. GEM generation was followed by a GEM-reverse transcription incubation, a clean-up step and 11 cycles of cDNA amplification. The resulting cDNA was evaluated for quantity and quality using a Thermo Fisher Scientific Qubit 4.0 fluorometer with the Qubit dsDNA HS Assay Kit (Thermo Fisher Scientific, Q32851) and an Advanced Analytical Fragment Analyzer System using a Fragment Analyzer NGS Fragment Kit (Agilent, DNF-473), respectively. Thereafter, 3’ gene expression libraries were constructed using a sample index PCR step of 14-15 cycles. The generated cDNA libraries were tested for quantity and size using fluorometry and capillary electrophoresis as described above. The cDNA libraries were pooled and sequenced with a loading concentration of 300 pM, paired end and single indexed, on an illumina NovaSeq 6000 sequencer using a NovaSeq 6000 S1 Reagent Kit v1.5 (100 cycles; illumina, 20028319) and a S4 Reagent Kit v1.5 (200 cycles; illumina, 20028313). The read set-up was as follows: read 1: 28 cycles, i7 index: 8 cycles, i5: 0 cycles and read 2: 91 cycles. The quality of the sequencing runs was assessed using illumina Sequencing Analysis Viewer (illumina version 2.4.7) and all base call files were demultiplexed and converted into FASTQ files using illumina bcl2fastq conversion software v2.20. At least 50,000 reads/cell were generated for each sample. All steps post cell-suspension preparation were performed at the Next Generation Sequencing Platform, University of Bern. Reads were aligned to the GRCh38 reference genome and quantified using CellRanger (version 6.0.0).

#### 6.1.2 Public datasets

When looking for datasets to include for this analysis, we excluded datasets with fewer than 20,000 genes as we wanted to ensure we captured as much of the transcriptome as possible. Table 2 shows where each dataset was obtained from.

**Table 2:** Summary of the publically available datasets included in this study and where they were obtained from.

| Dataset | Downloaded from | Comments |
| --- | --- | --- |
| Granja et al., 2019 [11] | Data was downloaded in the form of raw count matrices from GEO (GSE139369) and cell metadata was downloaded from the github repository ([GreenleafLab/MPAL-Single-Cell-2019](https://github.com/GreenleafLab/MPAL-Single-Cell-2019)) accompanying the publication. | Only the samples from healthy donors (BMMC_D1T1, BMMC_D1T2, CD34_D2T1, CD34_D3T1) were included in this analysis. |
| Oetjen et al., 2018 [12] | Data was downloaded in the form of raw count matrices from GEO (GSE120221). |  |
| Petti et al., 2019 [13] | Data was downloaded in the form of raw count matrices from the zenodo link accompanying the publication (https://doi.org/10.5281/zenodo.3345981 expression_matrices.tar file). |  |
| van Galen et al., 2019 [14] | Data was downloaded in the form of raw count matrices from GEO (GSE116256). | The 2 samples from AML cell lines (OCI-AML3 and MUTZ3) were excluded from this analysis. |

### 6.2 Analysis

#### 6.2.1 Pre-processing and normalisation

Datasets were analysed using a uniform pre-processing pipeline to ensure consistency. In the case where cell quality control needed to be performed, this was done using the ScanPy python module [64] according to current best-practices [65]. Counts were normalised using the `computeSumFactors` function from the scran R package [66]. The SingleR R package [67] was used to predict cell type labels for each dataset using the Granja et al., 2019[11] dataset as a reference. This dataset was chosen as the reference dataset as it is a CITE-seq dataset with expression values for several cell surface proteins and so the cell type labels should be reliable. Doublets were identified using Scrublet [68] and removed. All datasets were then merged into a single matrix for batch correction.

#### 6.2.2 Batch integration

Batch correction was performed using scArches [20] (2000 highly variable genes, 20 latent dimensions) and a nearest neighbour graph was constructed from the latent dimensions (n neighbours = 15) to generate a UMAP graph. A NextFlow [69] pipeline was developed to test different values for number of highly variable genes, latent dimensions and neighbours, to determine which values would yield the optimal integration results using the scPOP R package [70] to evaluate the results on the basis of Adjusted Rand Index (ARI), Normalized Mutual Information, Silhouette Width and Local Inverse Simpson Index (LISI). The results are shown in Supplementary Table 2.

#### 6.2.3 Cell type proportions

The `propeller` function from the `speckle` R package was used to test whether there are statistically significant changes in cell type proportions between the different conditions [21]. Samples that were sorted prior to sequencing (n = 6) were excluded from this analysis.

#### 6.2.4 Differential gene expression

For each cell type, differentially expressed genes were identified by comparing gene expression in healthy samples to gene expression in AML (diagnosis) samples. To avoid calling too many false positives, the Libra R package [25] was used to implement a pseudobulk approach to calculate differential expression and genes with absolute average log fold change *≥* 1 and an adjusted p value *≤* 0.05 were labelled as differentially expressed. Significant DEGs for each cell type were used as input for a Gene Ontology enrichment analysis using the `clusterProfiler` R package [71].

#### 6.2.5 Cell-cell interactions

For the prediction of cell-cell interactions, healthy and diagnosis samples were isolated from the larger dataset and 1000 cells (or all cells if n *≤* 1000) of each cell type were subsampled. The liana R package [30] was used to run 5 different methods for predicting cell-cell interactions (cellphonedb (squidpy), cellchat, NATMI, iTALK and SingleCellSignalR) using the Omnipath database resource of ligand-receptor interactions [31]. Results were filtered to interactions where either HSC.MPP cells were expressing either the ligand or receptor gene. To avoid calling false positive interactions, only interactions with `curation_effort` *≥* 3 were considered. The `liana_aggregate` function was used to generate an aggregate rank score for each interaction and interactions with a score *≤* 0.05 were considered to be high-confidence interactions. The `OmnipathR` R package was used to import functional annotations (KEGG and CellChat) for all genes involved in ligand-receptor interactions [31].

### 6.3 *In vitro* and *ex vivo* studies

#### 6.3.1 Reagents

2% sodium alginate was prepared by dissolving sodium alginate (Sigma) in phosphate buffered saline (PBS) and sterilised by heating to 80°C in a water bath for 15 mins [72]. Carboxyfluorescein succinimidyl ester (CFSE) (Biolegend) was dissolved in dimethyl sulfoxide (DMSO). Annexin V-APC was purchased from Immunotools. Anti-CD150-PE and anti-GRPC5C Alexa Fluor-405 antibodies were purchased from R&D Systems. Transforming growth factor beta-1 (TGFb-1) was obtained from Peprotech.

#### 6.3.2 Cell culture

KG1a cells were maintained at a density of 500,000 cells/ml in RPMI-1640 supplemented with GlutaMAX (Gibco, 2 mM), 10% HyClone fetal bovine serum (FBS, Thermo Fisher Scientific), penicillin (100 U/ml), streptomycin (100 *µ*g/ml) and sodium pyruvate (1 mM). Bone marrow mesenchymal stromal cells (BMSCs; hTERT immortalised primary BMSCs from a healthy donor) were cultured at a density of 50,000 cells/ml in *α*MEM (Sigma-Aldrich) supplemented with 10% HyClone FBS, penicillin (100 U/ml)/streptomycin (100 *µ*g/ml), sodium pyruvate (1 mM) and GlutaMAX (2 mM).

Primary AML blast samples. Mononuclear cell fractions generated from bone marrow aspirates were obtained from Blood Cancer Biobank Ireland. All patients provided written informed consent.

For BMSC-AML co-cultures, BMSCs were resuspended in 2% alginate solution at 500,000 cells/ml and aliquoted in 24-well plates (200 *µ*l/well). The alginate was crosslinked by addition of 100 mM CaCl2 for 10 seconds, forming a layer of BMSCs trapped in the alginate scaffold. Excess CaCl2 was removed by three washes with DPBS (Gibco). KG1a cells and primary AML blasts at 5×106 cells/ml density were labelled with the long-term cell tracker, carboxyfluorescein diacetate succinimidyl ester (CFSE, Biolegend, 2.5 *µ*M)) as per manufacturers instructions. The cells were resuspended in IMDM growth medium (supplemented with 10% HyClone fetal bovine serum (FBS, Thermo Fisher Scientific), penicillin (100 U/ml), streptomycin (100 *µ*g/ml) and sodium pyruvate (1 mM)) and seeded on top of the alginate-trapped BMSCs with or without TGFb1 (5 ng/ml) at 50,000 cells/ml for KG1a cells and at 500,000 cells/ml for primary AML samples.

#### 6.3.3 Immunofluorescence, cell cycle and viability analysis

Dye retention was analysed by detecting CFSE fluorescence intensity in the live cell fraction. Viability was determined using Annexin V-APC staining. KG1a cells previously labelled with CFSE were collected from single cultures by pipetting. From co-cultures, in addition to pipetting off the suspension cells, 200 *µ*l of 1 mM ethylenediaminetetraacetic acid (EDTA) was also added to loosen the alginate. The cells were pelleted and resuspended in Annexin V buffer (10 mM HEPES/NaOH, pH 7.5, 140 mM NaCl, 2.5 mM CaCl2) containing 1.5 *µ*l Annexin V-APC (ImmunoTools). The cells were incubated on ice in the dark for 15 min, followed by analysis by flow cytometry. The intensity of the CFSE signal was measured in the live cell fraction as a determinant of cell proliferation rate.

For immunofluorescence labelling, primary AML blasts were harvested and resuspended in 1% bovine serum albumin (BSA) in PBS and blocked for 10 mins after which the cells were collected and resuspended in 100 *µ*l 1% BSA/PBS containing 2 *µ*l of antibodies SLAMF1/CD150-PE (R&D Systems) or GRPC5C Alexa Fluor-405 (R&D Systems) and incubated on ice in the dark for 30 mins. After washing of unbound antibodies with 400 *µ*l of 1% BSA/PBS, the cells were resuspended in 200 *µ*l of 1% BSA/PBS for measurement.

## 7 Code and data availability

The scRNA-seq data generated as part of this study is available on GEO under the accession number GSEXXX. The code to reproduce the results is available at https://github.com/Sarah145/bone_marrow_analysis.

## 8 Supplementary materials

### 8.1 Supplementary Tables

- Supplementary Table 1: Clinical information for newly sequenced patients
- Supplementary Table 2: Sample information for all samples
- Supplementary Table 3: scPOP results
- Supplementary Table 4: DEG results
- Supplementary Table 5: Cell-cell interaction results

### 8.2 Supplementary Figures

**Figure 1:**
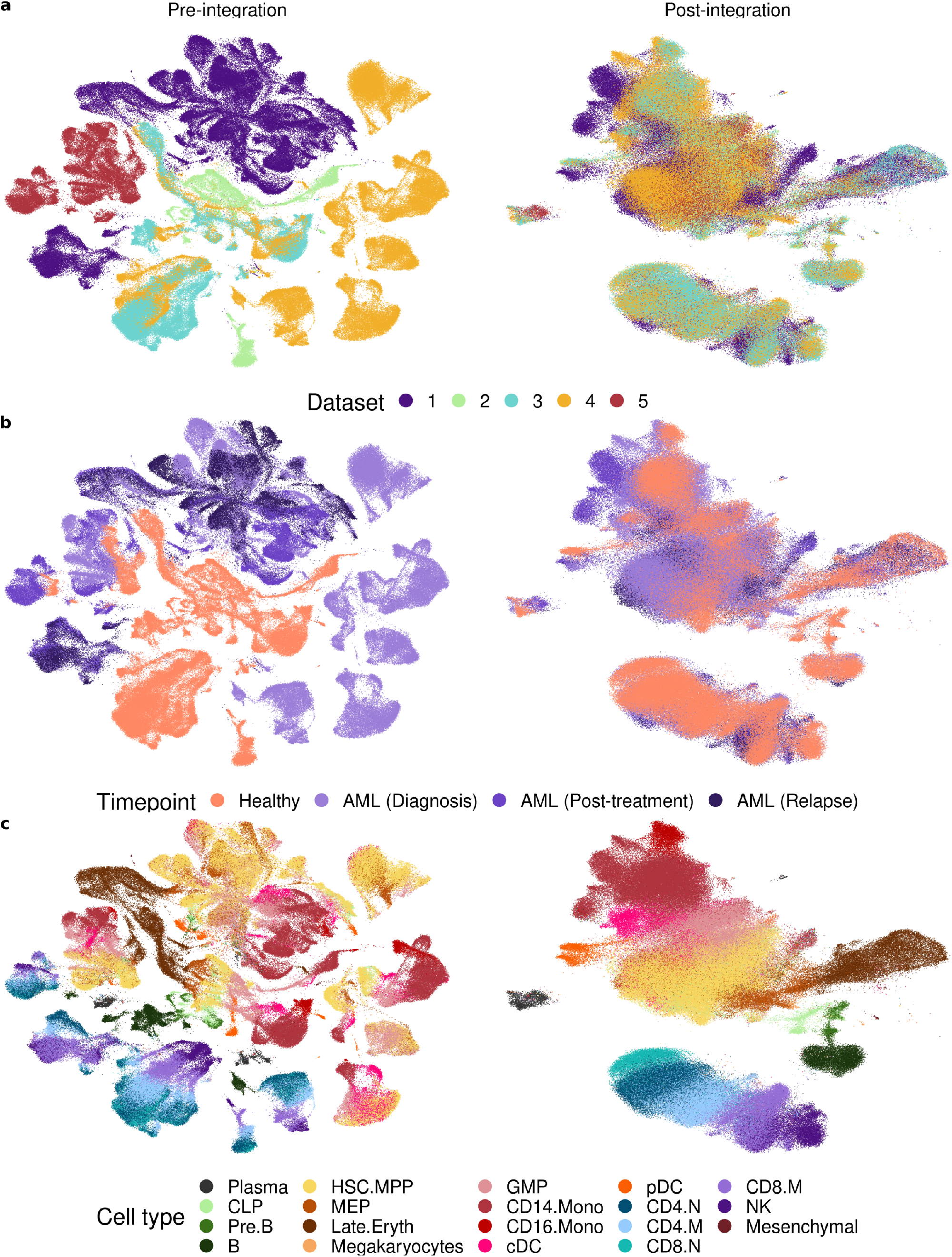
UMAP graphs pre-/post-integration.

**Figure 2:**
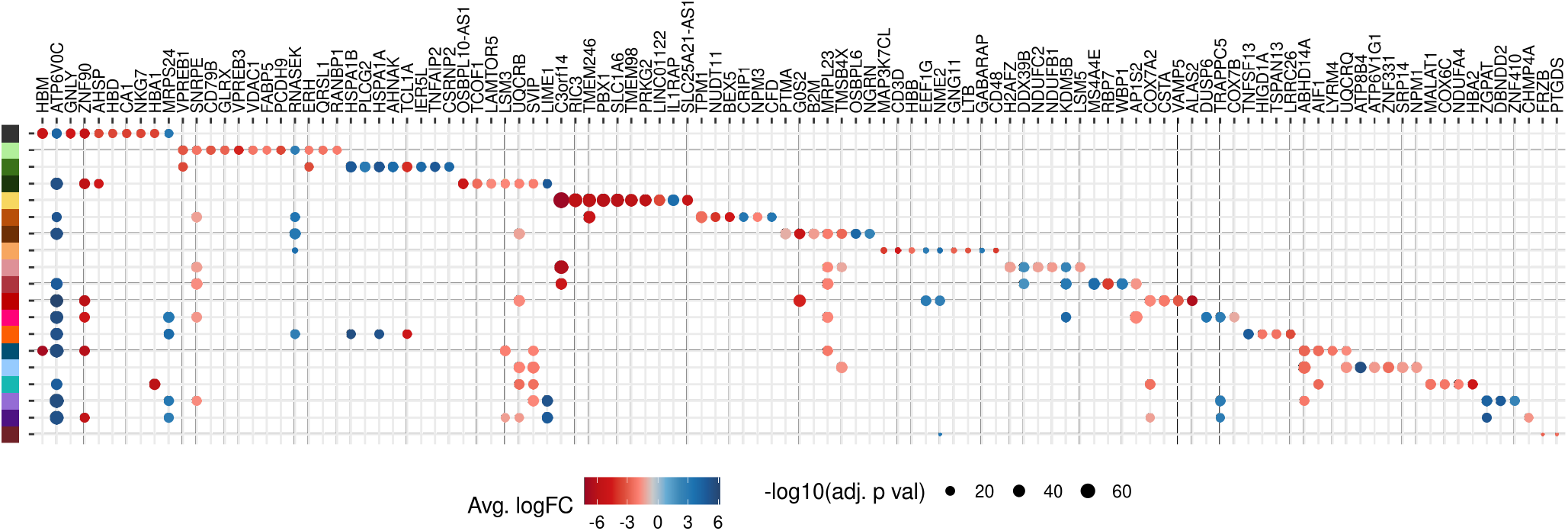
Top 10 DEGs for each cell type.

**Figure 3:**
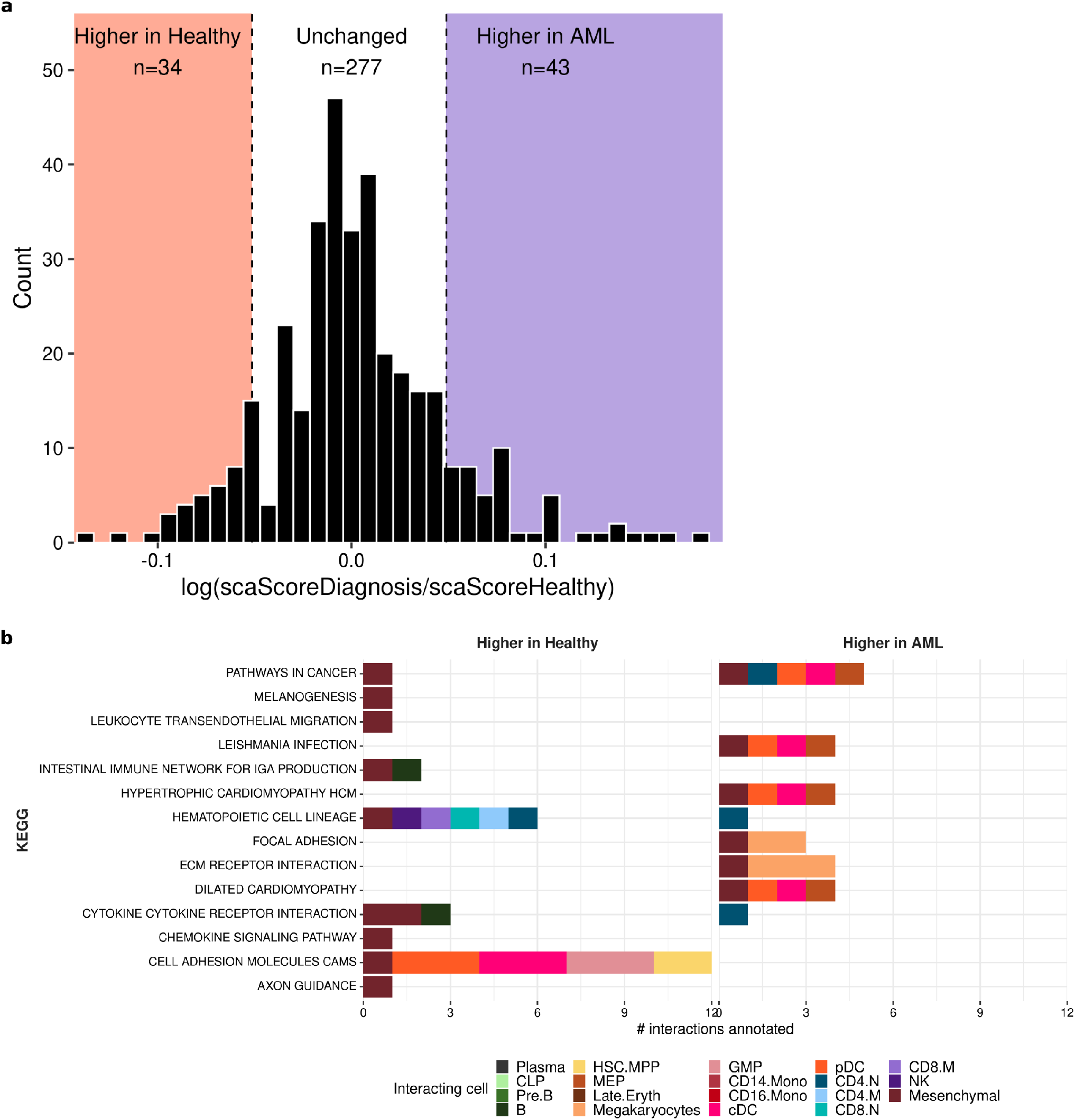
Interactions that were higher in healthy/AML.

**Figure 4:**
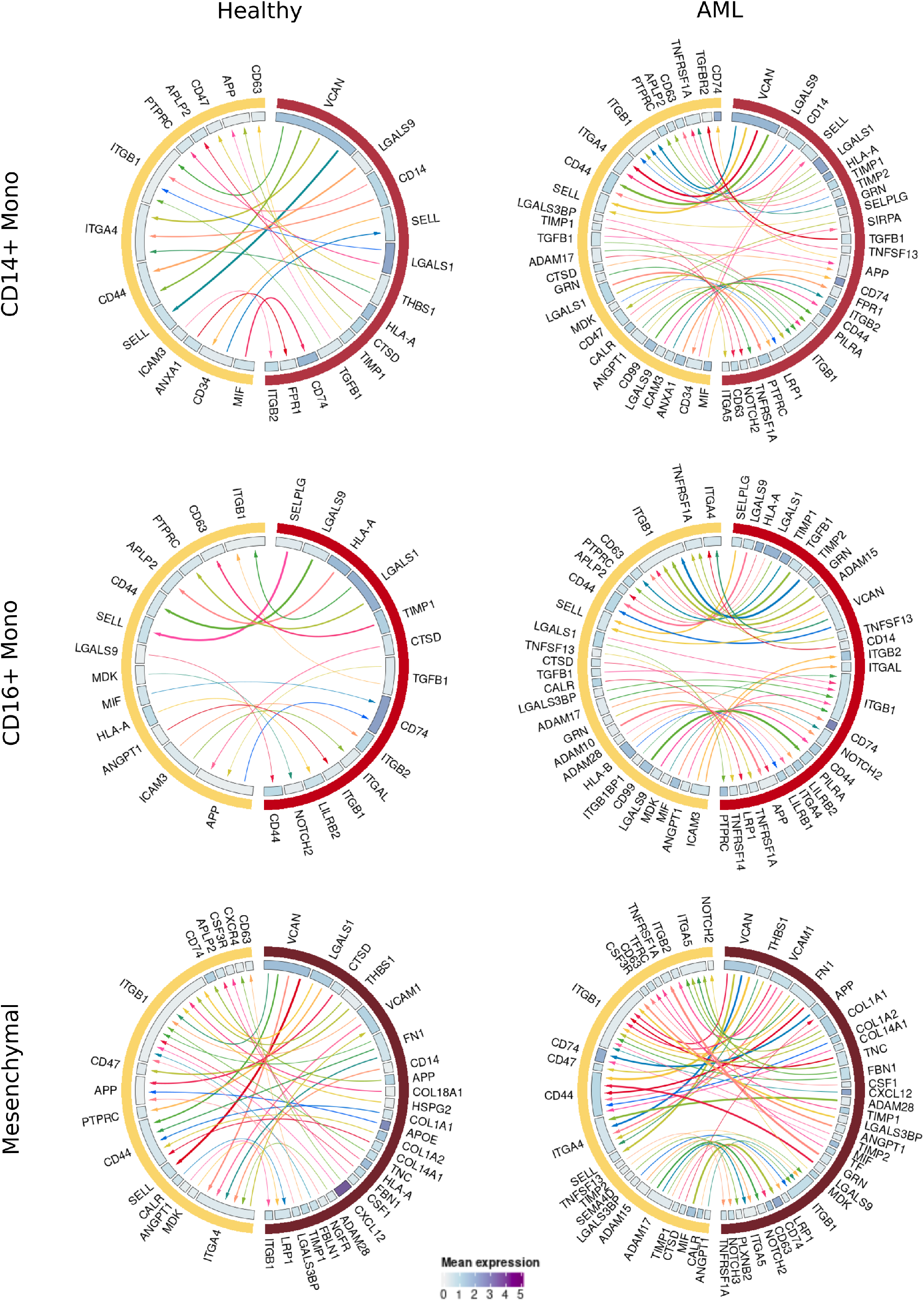
Interactions between HSC.MPPs and CD14.Mono, CD16.Mono and Mesenchymal cells.

